# Spectroscopic insights into the mechanism of anammox hydrazine synthase

**DOI:** 10.1101/2023.01.22.525020

**Authors:** Wouter Versantvoort, Rainer Hienerwadel, Christina Ferousi, Pieter van der Velden, Catherine Berthomieu, Laura van Niftrik, Frauke Baymann

## Abstract

Anaerobic ammonium oxidizing bacteria make a living oxidizing ammonium with nitrite as electron acceptor, intermediates nitric oxide and hydrazine, and end product dinitrogen gas. Hydrazine is a biologically unique free intermediate in this metabolism, and is produced by the enzyme hydrazine synthase. Crystallization of ‘*Candidatus* Kuenenia stuttgartiensis’ hydrazine synthase allowed for an initial hypothesis of its reaction mechanism. In this hypothesis, nitric oxide is first reduced to hydroxylamine after which hydroxylamine is condensed with ammonium to form hydrazine. Hydrazine synthase is a tetraheme cytochrome *c*, containing two proposed active site hemes (γI & αI) in the γ- and α-subunit, respectively, connected by an intra-enzymatic tunnel. Here we combined the data from electrochemistry-induced Fourier transform infrared (FTIR) spectroscopy, EPR and optical spectroscopy to shed light on the redox properties and protein dynamics of hydrazine synthase in the context of its reaction mechanism. Redox titrations revealed two low potential low spin hemes with midpoint potentials of ∼-360 mV and ∼-310 mV for heme αII and γII, respectively. Heme γI showed redox transitions in the range of 0 mV, consisting of both low spin and high spin characteristics in optical and EPR spectroscopy. Electrochemistry-induced FTIR spectroscopy indicated an aspartic acid ligating a OH^-^/H_2_O at the heme γI axial site as a possible candidate for involvement in this mixed spin characteristic. Furthermore, EPR spectroscopy confirmed the ability of heme γI to bind NO in the reduced state. Heme αI exhibited a rhombic high spin signal, in line with its ligation by a proximal tyrosine observed in the crystal structure. Redox titrations down to −610 mV nor addition of dithionite resulted in the reduction of heme αI, indicating a very low midpoint potential for this heme. *In vivo* chemistry at this heme αI, the candidate for the comproportionation of hydroxylamine and ammonium, is thus likely to be initiated solely on the oxidized heme, in contrast to previously reported DFT calculations. The reduction potentials of the γ-subunit hemes were in line with the proposed electron transfer of heme γII to heme γI for the reduction of NO to hydroxylamine (E^0^’ = − 30 mV).

## Introduction

Anaerobic ammonium oxidizing (anammox) bacteria are chemolithoautotrophic microorganisms that make a living by converting ammonium and nitrite to dinitrogen gas, with hydrazine as a unique free intermediate (1-3). Anammox bacteria are ubiquitous in both natural and engineered ecosystems, where they contribute significantly to the release of fixed nitrogen from that environment. It is estimated that approximately 50% of the nitrogen loss from the ocean is due to anammox activity (4, 5). Furthermore, they are successfully applied in over a 100 wastewater treatment plants worldwide (6-9).

The catabolic reactions of anammox metabolism take place in a dedicated intracellular compartment termed the anammoxosome (1, 2, 10). The current working hypothesis is that nitrite is initially reduced to nitric oxide by nitrite reductase (2, 11). This nitric oxide is subsequently condensed with ammonium to form hydrazine by hydrazine synthase (3). Hydrazine is then oxidized to dinitrogen gas by hydrazine dehydrogenase, releasing four low potential electrons (12). These electrons are shuttled through a membrane-bound respiratory chain, where they contribute to the maintenance of the proton motive force required for ATP synthesis, before they return into the anammoxosome to resume the first two steps of the anammox reaction cycle (2, 13).

The formation of hydrazine as a free metabolic intermediate is unique to anammox bacteria as is the enzyme responsible for its production, hydrazine synthase. The purification and crystallization of hydrazine synthase directly from native ‘*Candidatus* Kuenenia stuttgartiensis’, the type strain for anammox bacteria, allowed for an initial hypothesis of a two-step reaction mechanism. Hydrazine synthase crystallized as a dimer of the HZSα (kuste2861), HZSβ (kuste2860) and HZSγ (kuste2859) heterotrimer (**Figure 1**). The γ-subunit contains two *c*-type cytochromes and shows homology to the family of MauG and bacterial cytochrome *c* peroxidases (bCcP). Bis-His ligated heme γII (**Figure 1&2B**) is surface exposed and proposed to accept electrons from a cytochrome *c* partner, which it shuttles to heme γI (**Figure 1&2A**). Heme γI is covalently bound by three cysteines and is coordinated by a histidine and hydroxide. It is proposed to catalyze the three electron reduction of nitric oxide to hydroxylamine. Hydroxylamine then diffuses through an intra-enzymatic tunnel to the second active site, heme αI (**Figure 1&2C**). Ammonium diffuses to heme αI through a second, minor tunnel where it is condensed with hydroxylamine to from hydrazine. Strikingly, the histidine (His587) from the CxxCH binding motif of heme αI is coordinating a zinc residue and the heme has a tyrosine (Tyr591) as proximal ligand, reminiscent of catalases. Its protein environment, however is not homologous to any other known structure. The α-subunit contains an additional bis-his ligated *c*-type heme αII (**Figure 1&2D**), for which no function is yet hypothesized as it is not within efficient electron transfer distance from other HZS hemes. This heme is embedded in a typical type I cytochrome *c* fold. The β-subunit consists of a β-propellor structure, but does not contain any cofactors. It contains a loop region near the intra-enzymatic tunnel, which might regulate access to the α-subunit and likely fulfils a structural role (14).

**Figure 1:**
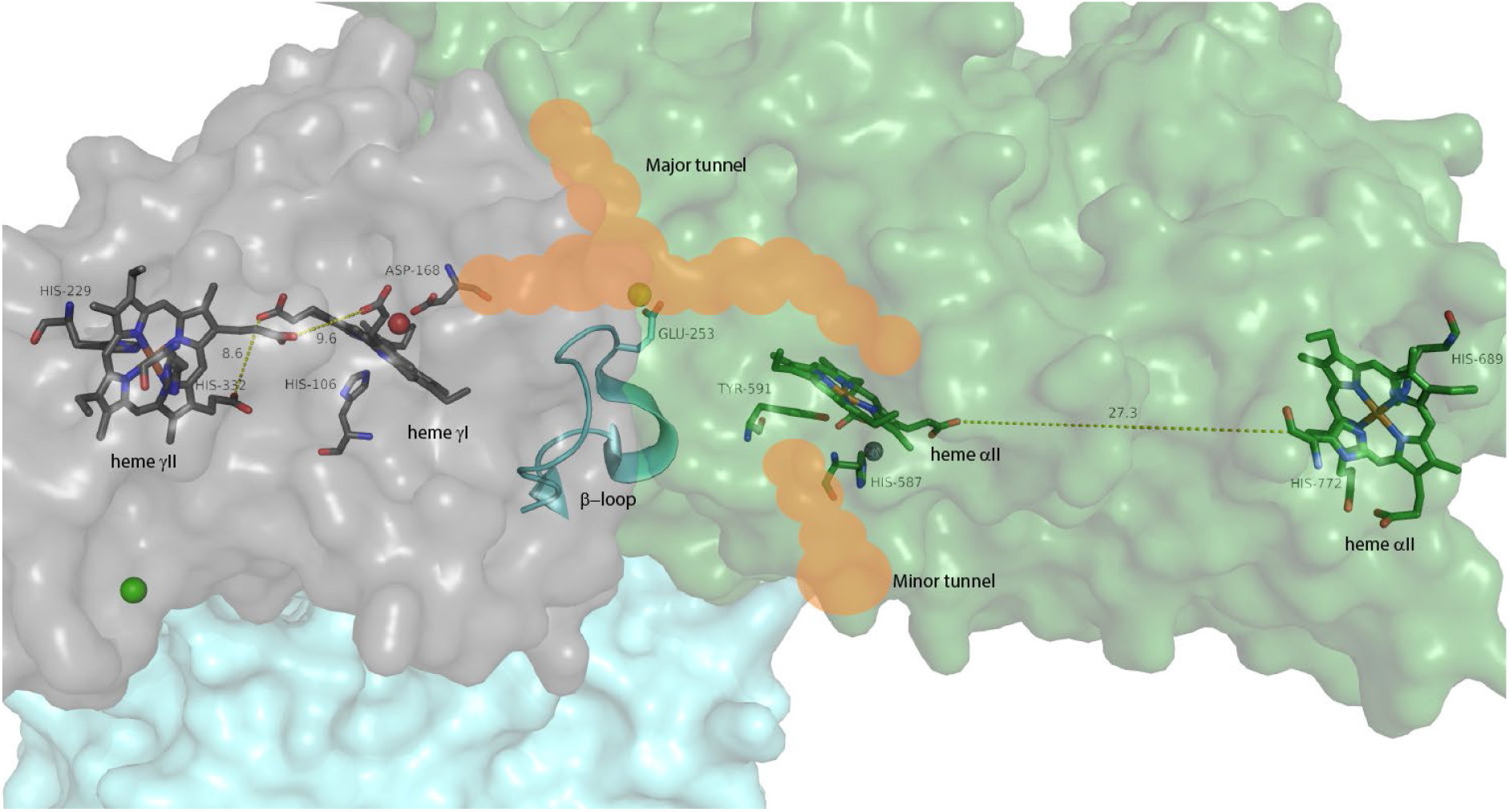
Crystal structure of the hydrazine synthase dimer from ‘Candidatus Kuenenia stuttgartiensis’, containing four c-type hemes per αβγ-monomer. The γ-subunit consists of a bis-his ligated heme yII, which is surface exposed near a cytochrome c binding site. Within 10 Å sits heme yI, coordinated by a proximal histidine and a hydroxide. The α-subunit contains a surface exposed bis-his coordinated heme αII and a 5-coordinated heme αI, with a tyrosine as proximal ligand. Heme αI and heme γI are connected by an intra-enzymatic tunnel. A loop in the β-subunit (245-260) approaches the tunnel and could regulate diffusion of molecules between the two hemes. (PDB:5C2V) (14).

Hydrazine synthesis is at the heart of the anammox metabolism and unique among biological energy conversion reactions. The molecular function of the enzyme proposed to catalyze this reaction is based on structural data and basic chemical knowledge. Experimental data to confirm the proposed mechanism (or come up with a different one) are missing. The redox properties of the enzyme cofactors and the dynamics of the protein at work can be assessed by electrochemistry coupled to various spectroscopic methods. Here we combined electrochemistry-induced Fourier transform infrared (FTIR) spectroscopy, EPR and optical spectroscopy to study the molecular mechanism of hydrazine synthase and compared the data to the proposed reaction mechanism.

## Materials and methods

### Protein purification

Hydrazine synthase purification was adapted from Kartal *et al*., 2011 (3) with some adjustments. All steps of the hydrazine synthase purification procedure were performed anaerobically. 1.5 L of anaerobic ‘*Candidatus* Kuenenia stuttgartiensis’ MBR1 bleed (O.D ∼ 1.2) (15) was centrifuged at 5000 x g, 15 min, 4 °C in a Allegra X-15R centrifuge (Swinging bucket rotor, Beckman Coulter). The cell pellet was resuspended in 10 ml 20 mM Tris pH 7.5 and passed once through a French press at 120 MPa. The resulting crude extract was subjected to ultracentrifugation at 162 000 x g, 50 min, 4 °C (Fixed angle Ti90 Rotor, Optima XE90, Beckman Coulter). The supernatant was loaded onto a 70 ml Q-sepharose column (XK 26/20, GE Healthcare/Cytiva) equilibrated with 20 mM Tris pH 7.5, connected to an Akta purifier (GE Healthcare/Cytiva) and run at a flow rate of 5 ml/min. Hydrazine synthase was eluted by a step to 20 mM Tris 200 mM, NaCl pH 7.5 and concentrated on a 50 kDa cutoff Amicon pressure filter unit. After buffer exchange to 20 mM KPi 100 mM NaCl pH 7, the sample was applied to a 10 ml ceramic hydroxyapatite Type-II 40 μm (Bio-rad) column (Omnifit, Diba Industries) equilibrated in the same buffer and run at 5 ml/min. First a step of 46 mM KPi, 100 mM NaCl pH 7 was applied to remove contaminants, after which hydrazine synthase was eluted with a step of 98 mM KPi, 100 mM NaCl pH 7. Hydrazine synthase was concentrated on a 100 kDa Vivaspin spinfilter (Sartorius) and stored at −20 °C in anaerobic serum bottles until use. Hydrazine synthase purity was checked using gel electrophoresis (16) and its identity confirmed using MALDI-ToF MS.

Hydrazine synthase α-subunit was purified by applying the 200 mM NaCl fraction from the Q-sepharose column on a 50 ml hydroxyapatite type II 40 μm (Bio-rad) column (XK 26/20, GE Healthcare/Cytiva) equilibrated with 20 mM KPi pH 7 at 5 ml/min. A step to 20 mM KPi 1 M NaCl was applied and the eluted sample concentrated and buffer exchanged to 20 mM KPi pH 7, before it was reapplied on the 30 ml hydroxyapatite column. A linear gradient of 0 to 1 M NaCl was applied and hydrazine synthase α-subunit eluted at a conductivity of 65 mS/cm. A 100 kDa cut-off spinfilter (Sartorius) was used to concentrate the sample and purity was assessed by gel electrophoresis.

### MALDI-ToF MS

Maldi-ToF MS was carried out according to Farhoud *et al*., 2005 (17). Briefly, bands of interest were excised from a SDS-PAGE gel and destained. After reduction and alkylation of cysteines with iodoacetamide, proteins were tryptically digested overnight. Peptides were extracted in 0.1 % trifluoroacetic acid in 50% acetonitrile, mixed in a 1:1 ratio with a 10 mg/ml 4-hydroxy-a-cyanocinnamic acid matrix solution and spotted on a MSP 96 stainless steel plate (Bruker). MALDI-ToF MS analysis was performed on a Microflex LT (Bruker) and peptides were mapped against an in-house ‘*Candidatus* Kuenenia stuttgartiensis’ MBR1 database using Biotools software, with carbamidomethylation as global and methionine oxidation as a variable modification, allowing for 1 missed cleavage site and a 0.3 Da mass tolerance.

### UV-Vis spectroscopy

UV-visible spectra were recorded aerobically on a Cary 60 spectrophotometer (Agilent) in a quartz cuvette (Hellma). After recording of as isolated spectra, samples were reduced using sodium dithionite. Origin 2020 (OriginLab Corp) was used to analyze the spectra.

### Optical redox titration

Optical redox titration was performed aerobically using an optically-transparent thin-layer electrochemical cell (OTTLE) (18) with CaF windows or a plexiglas version of similar design. Goldgrid (Buck Bee Mears) working electrodes were modified by boiling for 10 min in a 1 mg/ml PATS-3 solution. A platinum wire was used as a counter electrode and a Ag/AgCl_2_ wire in a 3 M KCl solution was used as a reference electrode, which was calibrated for SHE by titration of horse heart cytochrome *c*. The OTTLE was connected to a potentiostat and spectra were recorded on a customized Cary14 spectrophotometer from 700 to 375 nm at 10 °C. The sample consisted of ∼1 mM hydrazine synthase, 100 μM buffer Tris pH 8, 50 mM KCl and 40 μM (or 20 μM in the Plexiglas cell) mediators each (ferricyanide (E^0^′ = +430 mV); 1.4-benzoquinone (E^0^′ = +280 mV); 2,5-dimethyl-1,4-benzoquinone (E^0^′ = +180 mV); 1,2-naphthoquinone (E^0^′ = +145 mV); phenazine methosulfate (E^0^′ = +80 mV); 1,4-naphthoquinone (E^0^′ = +60 mV); phenazine ethosulfate (E^0^′ = +55 mV); 5-hydroxy-1,4-naphthoquinone (E^0^′ = +30 mV); 1,2-dimethyl-1,4-naphthoquinone (E^0^′ = 0 mV); 2,5-dihydroxy-p-benzoquinone (E^0^′ = −60 mV); 5,8-dihydroxy-1,4-naphthoquinone (E^0^′ = −145 mV), 9,10-anthraquinone (E^0^′ = −184 mV), 9,10-anthraquinone-2-sulfonate (E^0^′ = −225 mV); benzyl viologen (E^0^′ = −350 mV) and methyl viologen (E^0^′ = −440 mV/−772 mV).

Titrations were performed between +390 and −610 mV versus SHE in both reductive and oxidative directions in 30 mV steps, a 15 mV offset between reductive and oxidative cycles and 10 min equilibration time. The spectra were analyzed by global fitting using the mfit-nernst function of the QSoas software (19) to extract redox potentials and the corresponding difference spectra and plotted using Origin 2020. In addition, specific wavelengths were investigated as a function of potential (i.e. 406 nm minus 412 nm) using Origin 2020 and potentials were extracted by fitting to a sum of Nernst curves. The error in the obtained midpoint potentials was estimated at ± 20 mV.

### EPR spectroscopy

Hydrazine synthase samples were prepared in an anaerobic glovebox in 100 mM MOPS pH 7 and loaded into quartz EPR tubes, sealed with butyl rubber stopper and frozen inside the glovebox in a cold finger. Spectra were recorded on a Elexys E500 X-band EPR spectrometer fitted with a helium cryostat (Oxford instruments) for temperature control. Hydrazine synthase was reduced by addition of 1 equivalent of sodium dithionite per hydrazine synthase heme and reoxidized by addition of 1 equivalent of ferricyanide per hydrazine synthase heme. Parameters of the spectra are given in the figure legends. For experiments with nitric oxide, 45 μM hydrazine synthase was incubated in capped serum bottles containing a 5 % NO headspace (95 μM in solution) in the anaerobic tent.

### Electrochemically-induced Fourier transformed infrared difference spectroscopy

Sample preparation for FTIR experiments was identical to that for optical redox titration and before the sample was installed in the FTIR spectrophotometer, integrity was verified by taking a few optical spectra at different potentials. Then the cell was mounted in a Bruker Tensor 27 spectrometer, connected to a cooling system set at 10 °C and left for 2 hours to remove water vapor. Difference spectra were recorded in a high potential range (+390 to −210 mV) and low potential range (−210 to − 560 mV). At each potential the sample was equilibrated for 5 minutes before 300 scans were acquired and averaged. Several cycles were collected (>20) for each potential step and individual difference spectra were manually assessed before data from each cycle was averaged using the OPUS 7.5 software. Baseline drifts and sample loss were corrected for by subtracting the oxidized minus reduced from the reduced minus oxidized spectra.

### Structural analysis

Hydrazine synthase structural data (PDB: 5c2v) was analyzed using Pymol software (Schrödinger, LLC).

## Results & Discussion

### UV-Vis spectroscopy

Purified ‘*Candidatus* Kuenenia stuttgartiensis’ hydrazine synthase exhibited a ferric heme c UV-Vis spectrum in the “as isolated state” (**Figure 3, black line**) with characteristics for five- and six-ligated hemes as a Soret band maximum at 405.5 nm, a broad shoulder in the Q-band region between 500 and 600 nm, and a charge transfer band around 620 nm. Upon addition of dithionite, two distinct spectra appeared in a time-dependent manner. Immediately after dithionite addition, part of the Soret band shifted to 419 nm with a concomitant appearance of a split alpha band with maxima at

**Figure 2:**
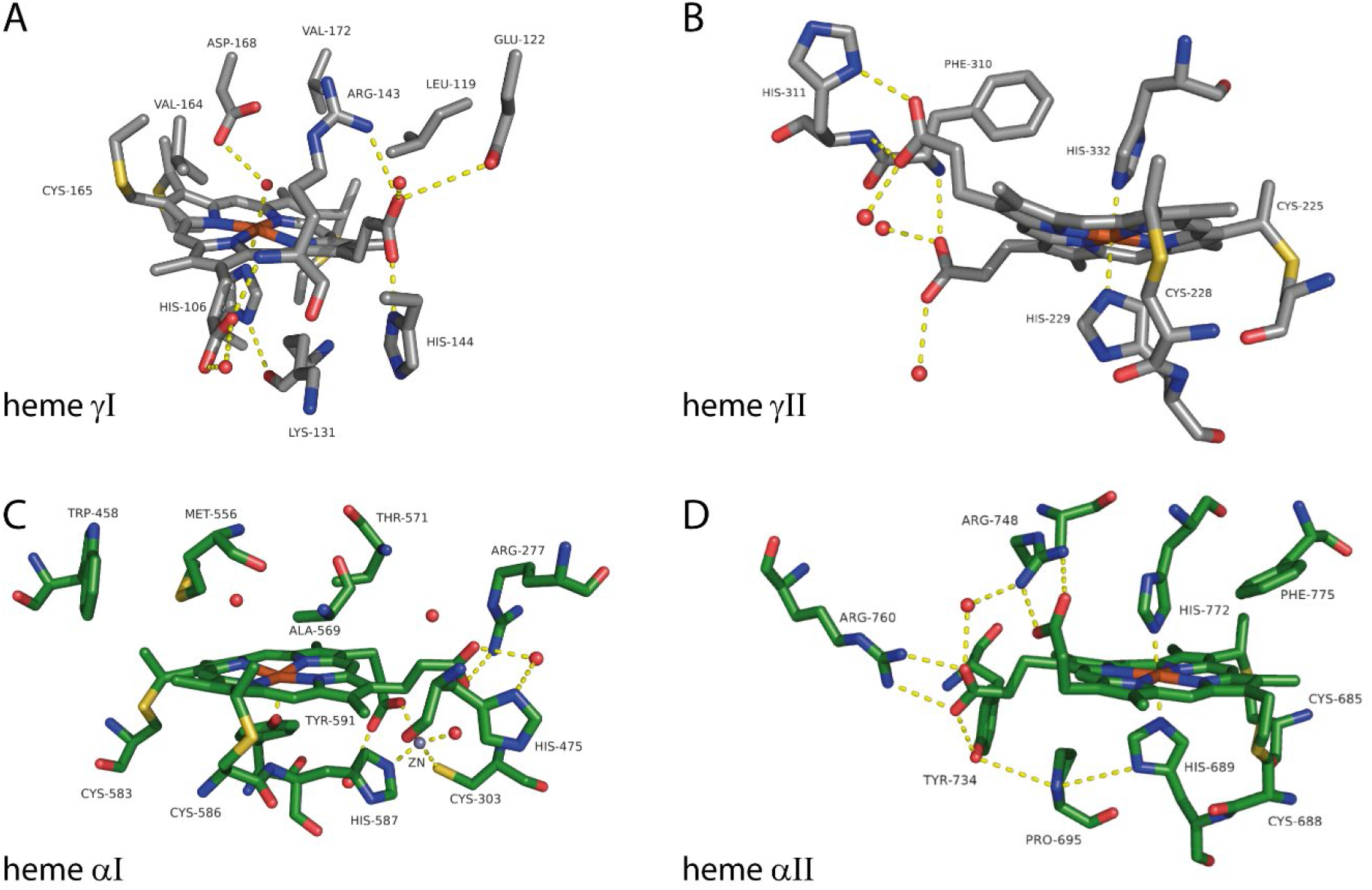
Close-up of view hemes γI **(A)**, γII **(B)**, αI **(C)** and αII **(D)** of ‘Candidatus Kuenenia stuttgartiensis’ hydrazine synthase and their immediate environment (PDB:5C2V) (14).

**Figure 3:**
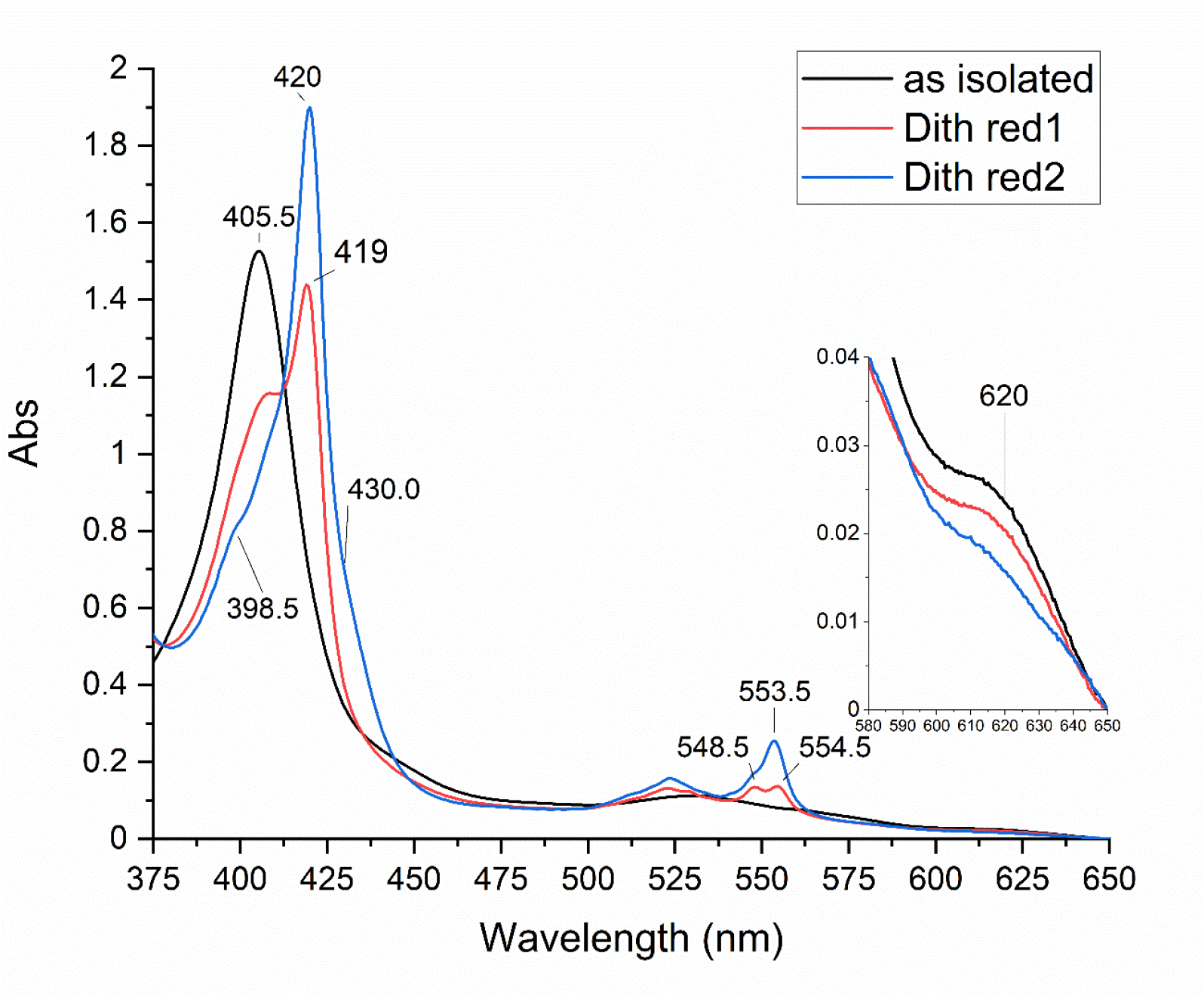
UV-Vis spectra of ‘Candidatus Kuenenia stuttgartiensis’ hydrazine synthase “as isolated” (black) and reduced with dithionite. Immediately after dithionite addition (red) part of the hemes got reduced, resulting in the appearance of two alpha bands with maxima at 548.5 and 554.5 nm and a shift in the Soret region from 405.5 nm to 419 nm. After incubating with dithionite for 5 minutes (blue) the alpha region showed a peak with a maximum at 553.5 and the Soret peak was shifted to 420 nm. A hint of a shoulder at 398.5 nm was visible in this dithionite reduced sample. Inset shows the presence of a charge transfer band at 620 nm, typical for high spin hemes. This band did not decrease immediately upon dithionite addition (red), but decreased after 5 min incubation of the sample with dithionite (blue).

548.5 nm and 554.5 nm (**Figure 3, red line**). No intensity decrease in the charge transfer band was detected. After incubating with dithionite for 5 minutes, the contribution at 405.5 nm was further decreased and the Soret maximum was shifted to 420 nm. This Soret band contained two shoulders at 398.5 and 430 nm, respectively. Furthermore an alpha band maximum, typical for a reduced low spin heme was observed at 553 nm, overlaying the split alpha band (**Figure 3, blue line**). Additionally, the charge transfer band at 620 nm drastically decreased in intensity, which together with the appearance of the 430 nm shoulder indicated the reduction of a high spin heme. The reduced minus oxidized difference spectra (**Figure 4**) calculated for the immediate reduction step, i.e. split-peak minus “as isolated” (**Figure 4, red line**) and for the second reduction step, i.e. reduced minus split-peak (**Figure 4, blue line**), corroborated the appearance of three distinct spectral components. The split-peak minus “as isolated” spectrum showed a shift of the Soret band from 404 nm to 420 nm and the appearance of the split alpha band with maxima at 549 and 554.5 nm. The difference spectrum of the 5 minutes incubated sample minus the split-peak spectrum showed a further decrease of the Soret at 403.5 nm with an increase of absorbance at 421.5 nm and a narrow alpha band at 553 nm. Additionally, a clear shoulder appeared at 429 nm and the 620 nm signal disappeared.

**Figure 4:**
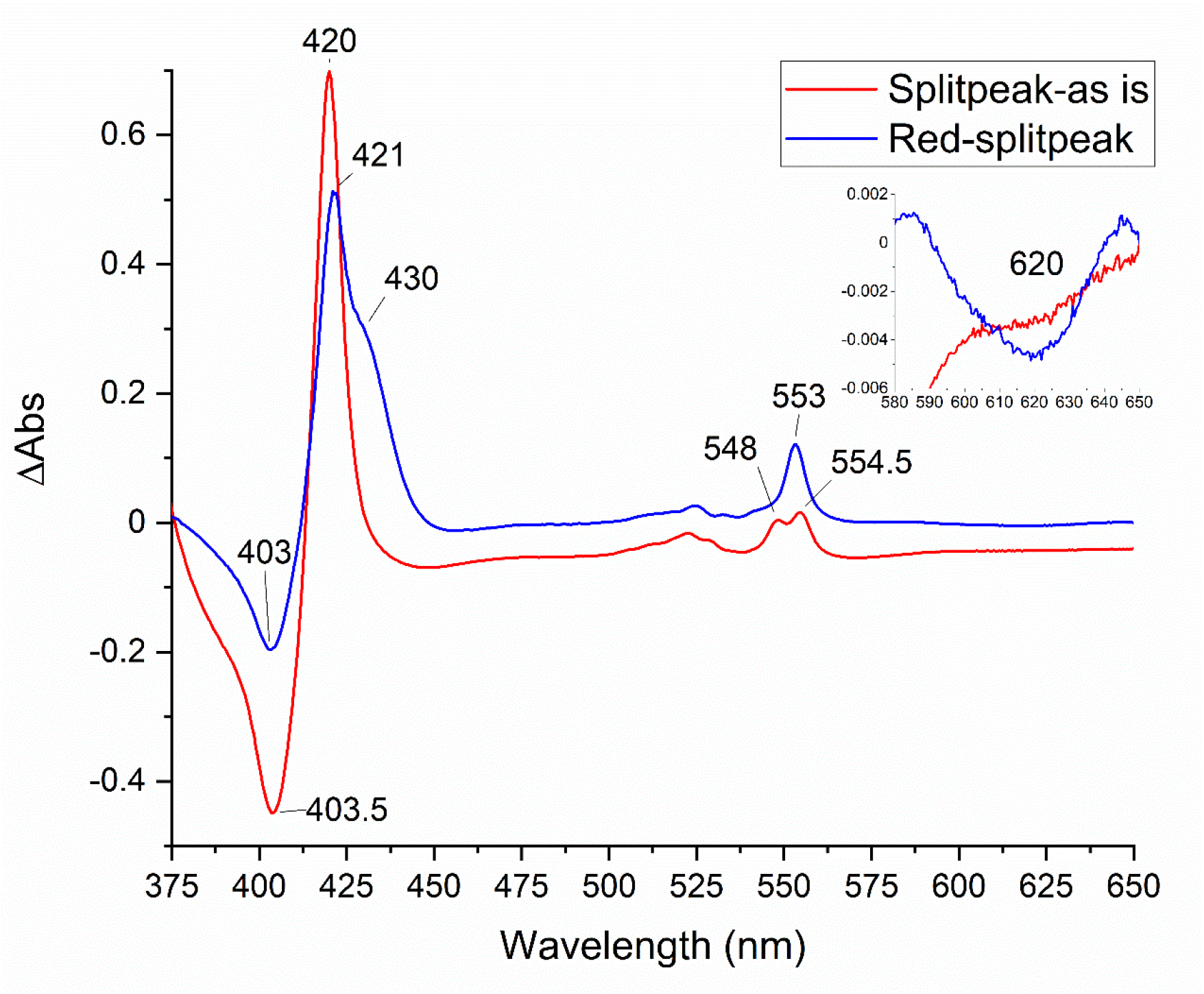
UV-Vis difference spectra of ‘Candidatus Kuenenia stuttgartiensis’ hydrazine synthase. The difference spectrum of hydrazine synthase immediately after dithionite addition minus the “as isolated” spectrum (red), showed a shift in the Soret maximum from 403.5 to 420 nm and the appearance of the double peak in the alpha region with maxima at 548 and 554.5 nm (therefore named splitpeak spectrum). The difference spectrum of hydrazine synthase incubated for 5 min with dithionite minus the splitpeak spectrum (blue) showed the appearance of a peak in the Soret region at 421 nm with a shoulder at 430 nm and a peak at 553 nm. The inset shows a zoom of the 580 to 650 nm range, where the decrease in the intensity of the charge transfer band at 620 nm was visible in the red minus splitpeak spectrum (blue).

The UV-Vis spectra of the purified hydrazine synthase α-subunit showed a similar “as isolated” spectrum (**Figure 5, black line**) as the complete hydrazine synthase, with a Soret maximum at 405.5 nm, a broad shoulder in the Q-band region and a charge transfer band at 615 nm. Reduction of the α-subunit with dithionite resulted in a shift of the Soret band to 419.5 nm, with a shoulder at 398.5 nm, and the appearance of a split alpha band with maxima at 548 and 554.5 nm (**Figure 5, red line**). This spectrum was very similar to that obtained immediately after dithionite addition to the whole enzyme (**Figure 3, red line**) and indicated that these contributions stem from hemes in the α-subunit. Additional incubation of the α-subunit with dithionite did not result in the appearance of a shoulder at 430 nm nor an alpha band at 553 nm, nor the disappearance of a charge transfer band, as was observed for the whole enzyme, which indicated these later contributions stem from hemes of the γ-subunit.

**Figure 5:**
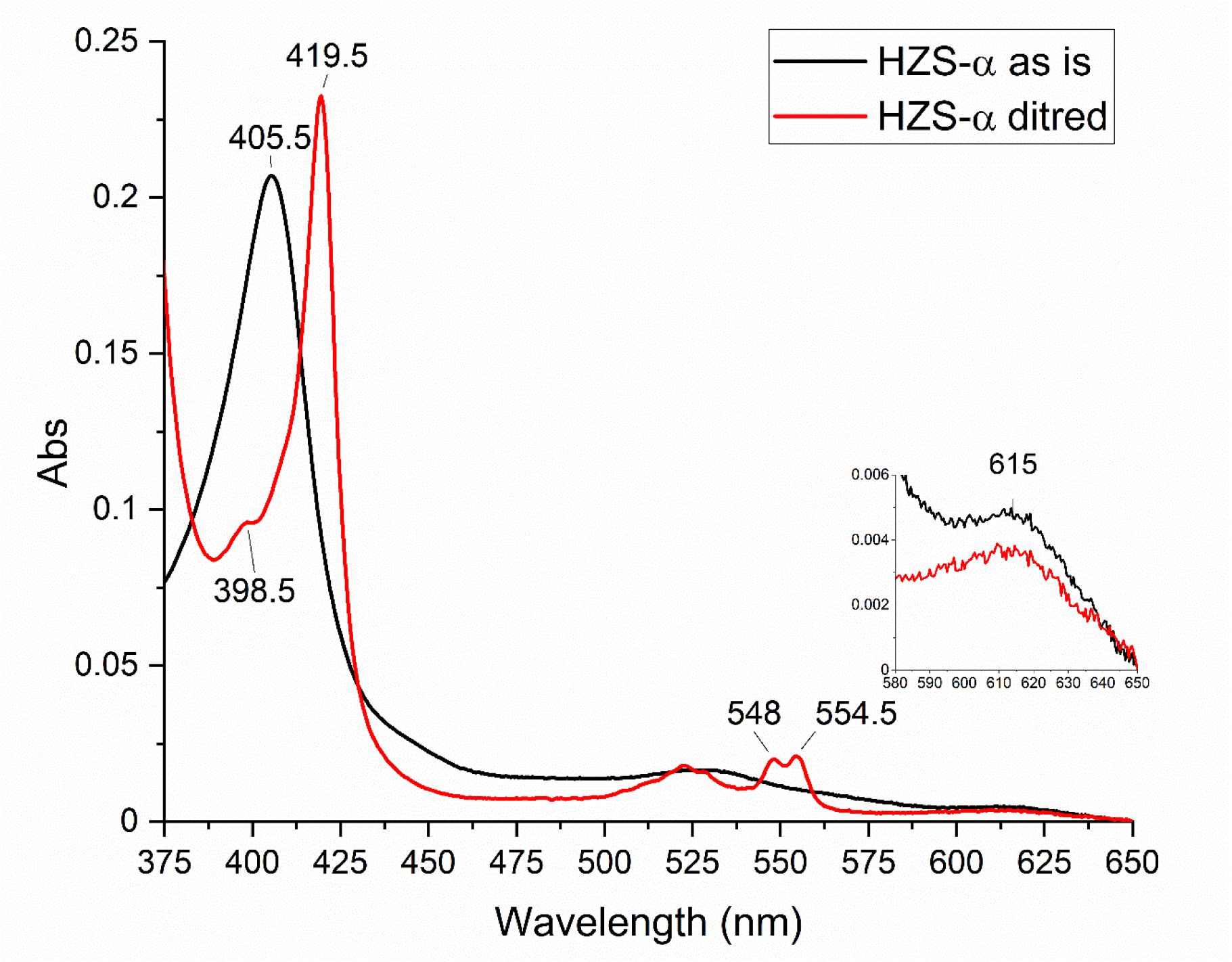
UV-Vis spectra of the isolated hydrazine synthase α-subunit from ‘Candidatus Kuenenia stuttgartiensis’ in the “as isolated” (black) and dithionite reduced (red) state. Upon dithionite reduction, two peaks appeared in the alpha region with maxima at 548 and 554.5 nm and the Soret maximum shifted to 419.5 nm. A minor peak at 398.5 nm became apparent in the dithionite reduced sample. The inset shows a zoom of the 580 to 650 nm region, where the presence of a charge transfer band at 615 nm was detected, which remained upon dithionite addition.

### EPR spectroscopy

The EPR spectrum of the “as isolated” anaerobic hydrazine synthase showed a rhombic signal (HSp1) with a g_z_ at 6.29 and g_y_ at 5.65, and an axial signal (HSp2) at g = 6 in the high spin region (**Figure 6, black line**). Two highly anisotropic low spin signals were observed, HALS1 with g_z_ at 3.08 and HALS2 with g_z_ 3.46, respectively. Additionally, a low spin ferric heme signal (LS1) is observed with a g_z_ at 2.57, g_y_ at 2.27 and g_x_ at 1.8. Lastly, a radical signal is observed at g ≈ 2. Due to the different angles between the histidines ligating the low spin hemes, 30° for heme αII (**Figure 2D**) and 40° for heme γII (**Figure 2B**), respectively, HALS1 could be attributed to heme αII and HALS2 to heme γII, like previously determined for aerobically purified hydrazine synthase (14). This aerobic preparation also showed similar signals in the high spin region, with both an axial and rhombic signal, but contained two additional low spin signals (14). These additional signals could have been induced due to oxygen exposure of the enzyme, as in our preparation hydrazine synthase was purified anaerobically. Reduction of hydrazine synthase with one equivalent of dithionite resulted in the disappearance of all low spin signals and the axial high spin signal, whereas the rhombic high spin signal remained. Further addition of dithionite did not change this. The rhombicity of the signal is characteristic for a tyrosine-ligated high spin heme and we therefore attributed this signal to heme αI. This heme αI, irreducible by dithionite, should give rise to the Soret band at 398.5 nm observed in the dithionite reduced spectra of the whole enzyme (**Figure 3, blue line)** and the isolated α-subunit (**Figure 5, red line)**. The reducible, axial high spin heme signal HSp2 then can be attributed to the histidine-ligated heme γI (**Figure 2A**). Its small signal size, when compared to the rhombic signal of heme αI showed that only part of the population of the γI heme was in a spin state that gave rise to signal in the g=6 region. Another population of the same heme may be at the origin of the LS1 signal. Interestingly, reoxidation of the sample with one equivalent of ferricyanide recovered all signals apart from LS1.

**Figure 6:**
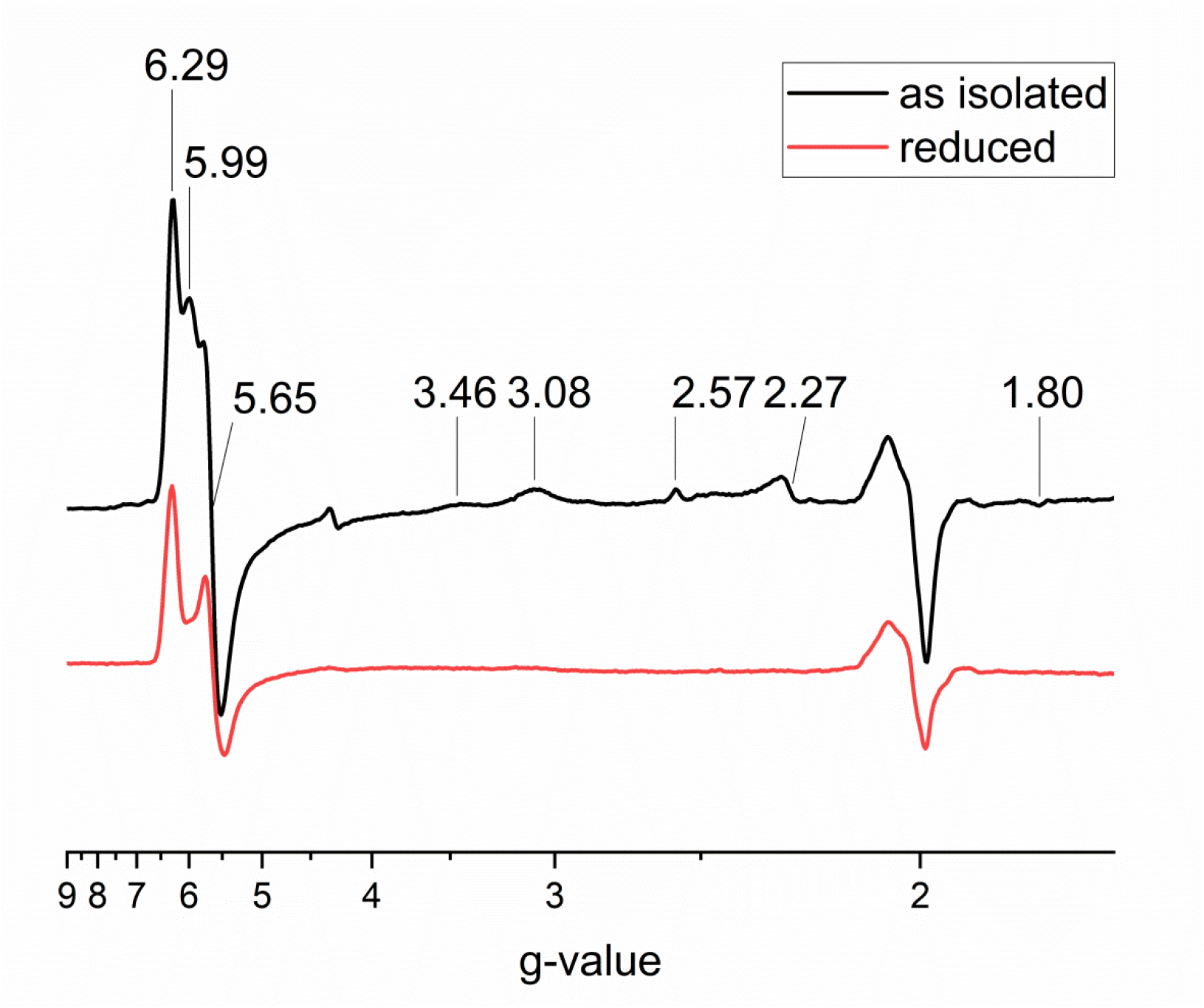
X-band EPR spectra of ‘Candidatus Kuenenia stuttgartiensis’ hydrazine synthase recorded at 25 K with 15 dB modulation amplitude. As isolated (**black**) anaerobic hydrazine synthase showed three signals in the high spin region with g-values of 6.29, 5.99 and 5.65, two highly anisotropic low spin signals with g values of 3.46 and 3.08, and one low spin signal with g-values of 2.57, 2.27 and 1.80. Lastly, a radical signal at g = 2 was observed. Reduction of hydrazine synthase with one equivalent of dithionite (**red**) results in the disappearance of all signals except for part of the high spin signal with g values of 6.29 and 5.65 and the signal around g =2.

To better resolve the signal at g ≈ 2 we recorded spectra on a narrow range with a modulation amplitude of 6 Gauss at higher temperature (60 K). These conditions resolved the spectral characteristics of a NO radical bound to a reduced heme. The three lines resulting from interaction of the radical spin with the nuclear spin of the nitrogen were spaced by 16 G, without further hyperfine splitting, which is characteristic for NO bound as a fifth ligand to a heme (**Figure 7, black**) (20). As NO is one of the substrates for hydrazine synthase, part of the enzyme might have been trapped in an NO-bound state upon cell disruption. Incubating reduced hydrazine synthase with NO resulted in an increased signal in this spectral region (**Figure 7, red**) and we estimated the spin concentration to about half the amount of hydrazine synthase present in the sample (28 μM NO-bound heme for 45 μM HZS).

**Figure 7:**
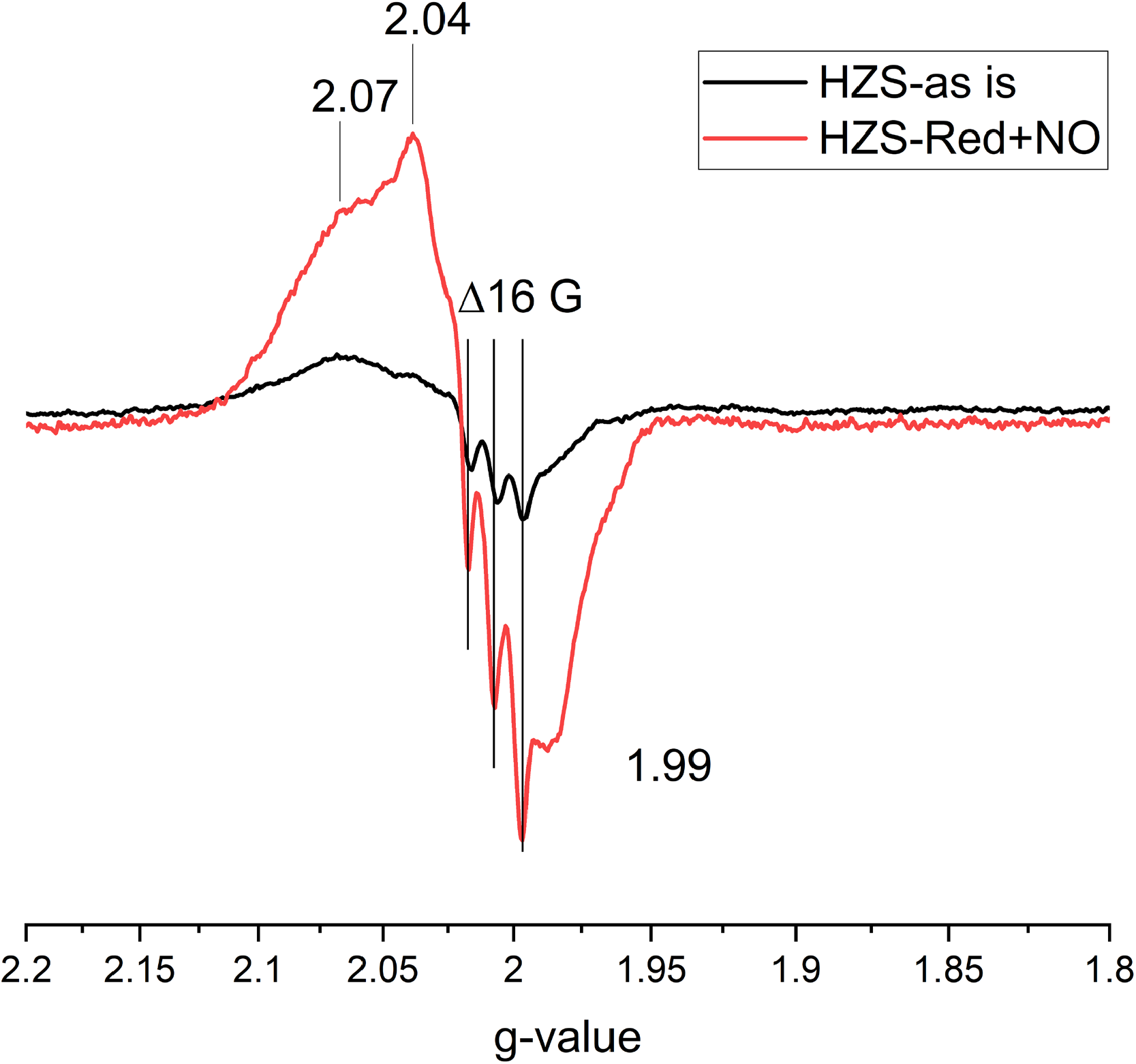
X-band EPR spectra of ‘Candidatus Kuenenia stuttgartiensis’ hydrazine synthase recorded at 60 K with a 15 dB modulation amplitude. “As isolated” hydrazine synthase (black) and dithionite reduced hydrazine synthase treated with ∼95 μM NO (red) both showed a radical signal with a 16 G hyperfine split of the g_y_-tensor. Spectra were normalized on protein concentration.

### Optical redox titration

An optical redox titration of hydrazine synthase at pH 8 (**Figure 8**) was used to determine the reduction potentials of the different *c*-type hemes. Upon reduction, the Soret band at 405.5 nm decreased and an initial absorbance increase at 430 nm was observed in combination with an alpha band at 554 nm and a decrease of the charge transfer band at 620 nm. From −200 mV onwards a Soret band with final maximum at 419.5 nm started appearing in combination with Q-bands at 523 and 553 nm. Absorbance changes at 406-412 nm were plotted against the ambient redox potential (**Figure 9)**. Two separate absorbance decreases indicated heme reduction occurring around 0 mV and −320 mV. The low potential wave was fully reversible in the oxidized direction, whereas the high potential wave was not. Overall 5% of the Soret intensity was lost upon reoxidation. A sum of three n=1 Nernst curves with redox midpoint potentials of −360 mV, −310 mV and −30 mV best described the data. Whereas the fitted line followed the data point in the low potential range a deviation from a n=1 nernstian behaviour was visible around 0 mV. Multiple redox cycles over different experiments showed reproducible, overlaying signals for the low potential part, but variability in the high potential region.

**Figure 8:**
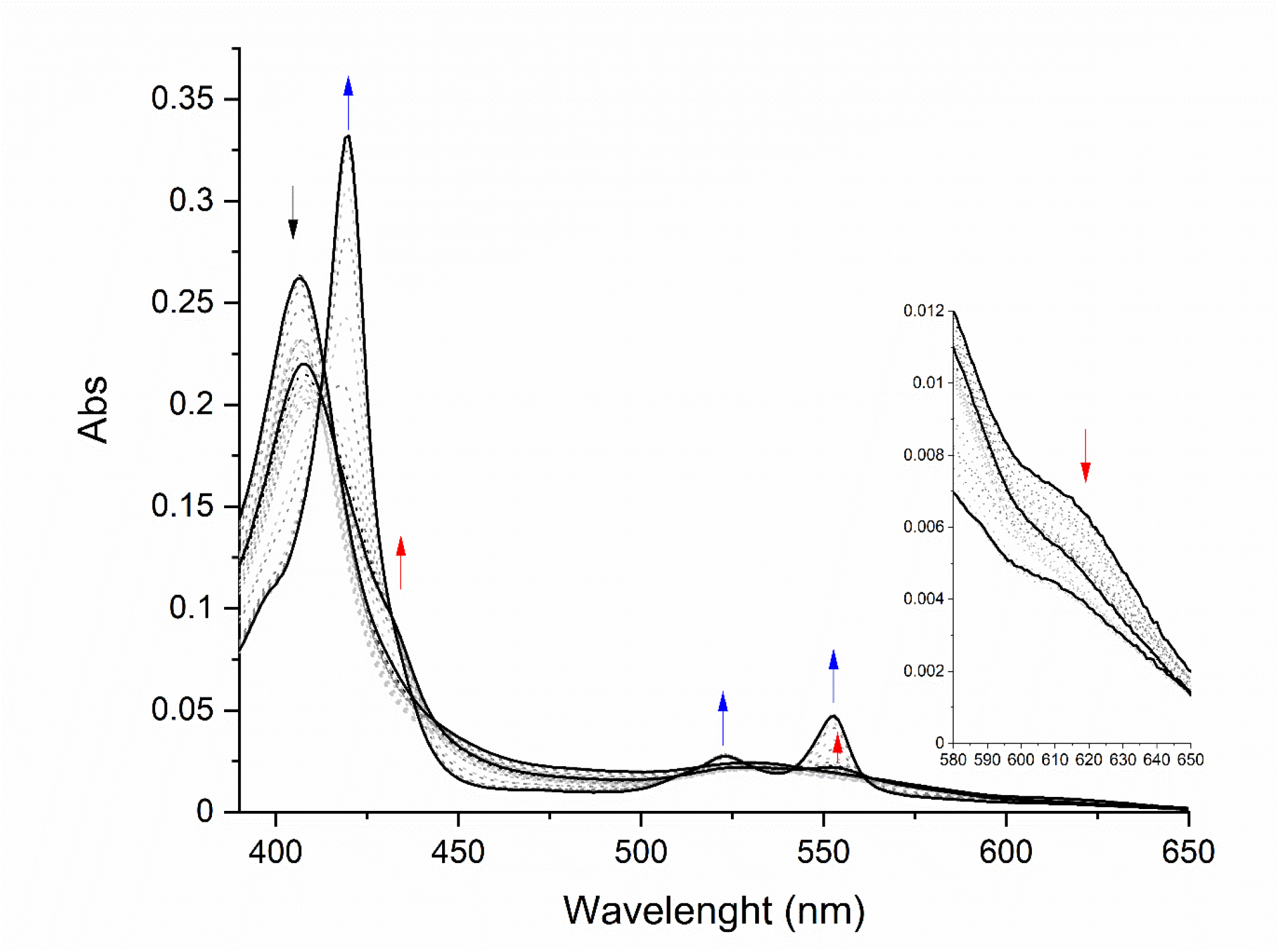
UV-Vis spectra of ‘Candidatus Kuenenia stuttgartiensis’ hydrazine synthase during the first reduction and subsequent oxidation cycle of a redox titration at pH 8. Upon reduction the Soret band at 405.5 nm and the charge transfer band at 620 nm decreased with a concomitant initial increase of a Soret band at 430 nm and an alpha-band at 554 nm (red arrows). Below −200 mV a Soret band with a final maximum of 419.5 nm started appearing in combination with the appearance of Q-bands at 523 and 553 nm, without further changes to the charge transfer band (blue arrows).

**Figure 9:**
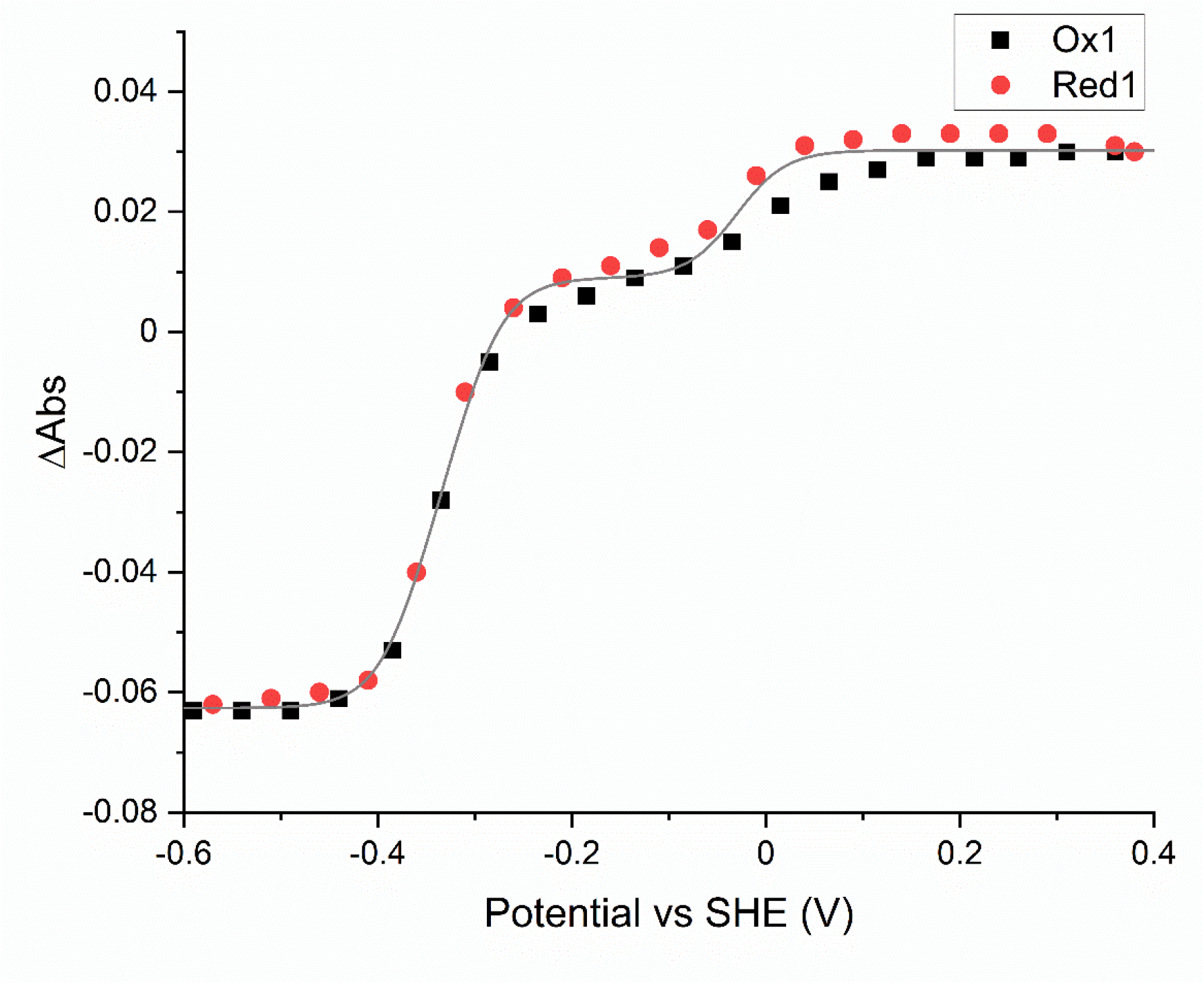
Amplitude changes of the absorbance difference between 406 nm and 412 nm as a function of the reduction potential. During the oxidation and reduction absorbance changes followed the same behavior in the low potential range, but not in the high potential range, where they deviated from an n=1 Nernst curve. The line represents a sum of three n=1 Nernst curves with redox midpoint potentials of −360, −310 and −30 mV.

Through a multicomponent Nernst fit, three distinct spectral components with reduction potentials of approximately −360 mV, −310 mV and −30 mV, respectively, could be extracted (**Figure 10**). The two low potential components showed reduced minus oxidized difference spectra characteristic of low spin hemes. The species with the lowest reduction potential (*E*_*m*_ ≈ −360 ± 20 mV), showed a split alpha band, indicating it corresponds to heme αII. The second low potential species (*E*_*m*_ ≈ −310 ± 20 mV) showed a Soret band at 420 nm and alpha band at 553 nm in the reduced minus oxidized difference spectrum, in agreement with the spectral changes ascribed to hemes in the γ-subunit, thus belonging to heme γII. The high potential spectrum (*E*_*m*_ ≈ −30mV in this experiment but a larger variability between redox cycles and experiments than for the low potential hemes was observed) had a broad Soret maximum at 429 nm and a weak alpha band at 554 nm, in addition to a negative charge transfer band at 620 nm. This indicated a mixed high spin/low spin contribution to this spectrum, which was in agreement with the attribution of the g = 6 and the LS1 EPR signals to heme yI.

**Figure 10:**
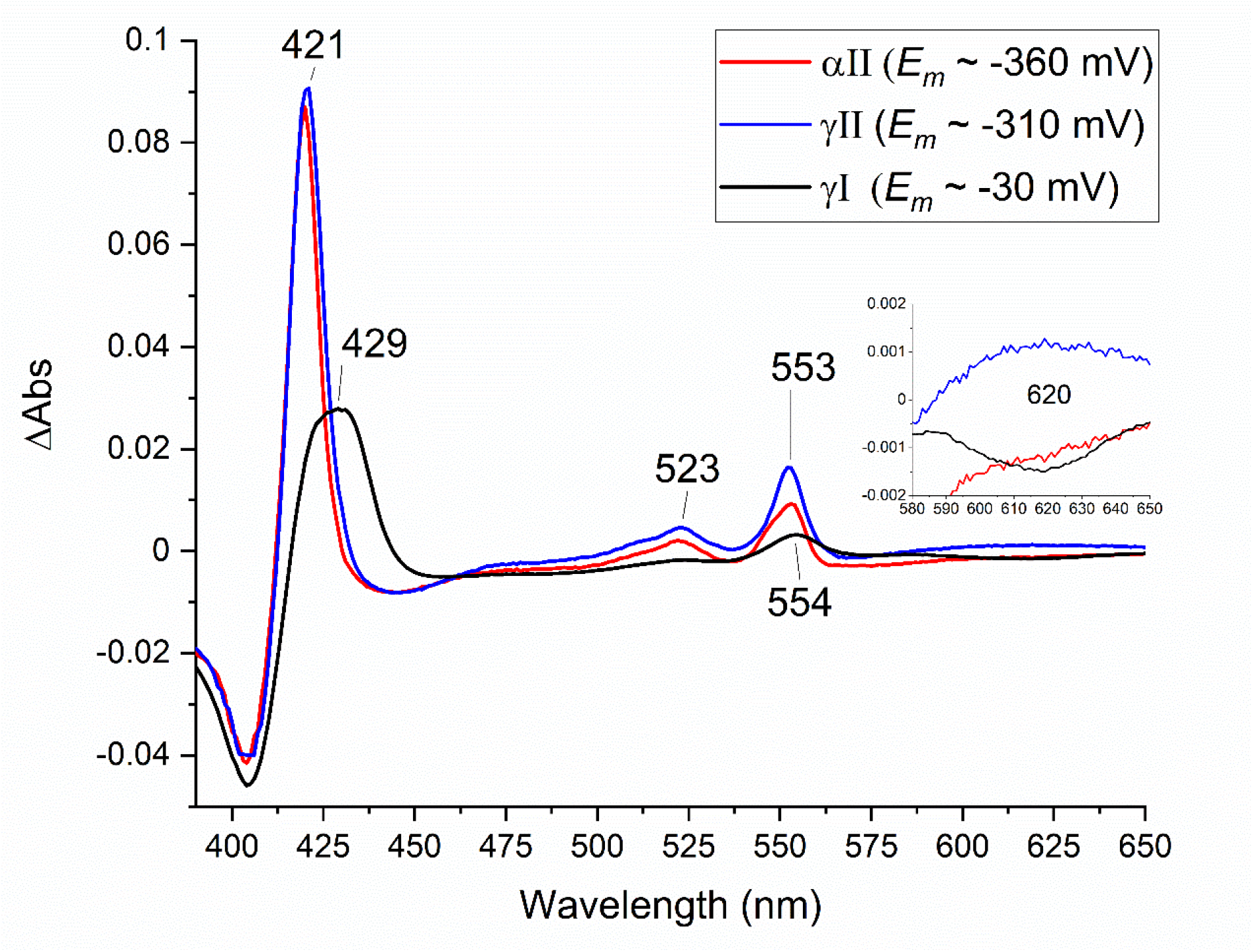
UV-Vis spectra of the 1^st^ reduction cycle of a redox titration of ‘Candidatus Kuenenia stuttgartiensis’ hydrazine synthase at pH 8 as extracted from the titration data by a global fit analysis (QSoas (19)) with three components.

### FTIR difference spectroscopy

The molecular changes associated with the reduction of the hemes were investigated using electrochemistry-induced FTIR difference spectroscopy. Heme modes as well as contributions from axial ligands, propionate groups, amino acid side-chains or the protein backbone are expected to contribute to the FTIR difference spectra. Reduced-minus-oxidized FTIR difference spectra were first recorded for the +390 mV to −210 mV range (**Figure 11A**) and for the −210 mV to −560 mV range (**Figure 11B**). According to the multicomponent Nernst fit, EPR and UV-Vis data, the FTIR difference spectrum reported in Figure 11A corresponded mainly to the reduction of the “high potential” heme γI. The FTIR spectrum reported in Figure 11B should correspond to contributions of both low potential species with *E*_*m*_ ≈ −360 mV and *E*_*m*_ ≈ −310 mV, assigned to hemes αII and γII, respectively. By splitting the −210 mV to −560 mV potential step in two separate ranges from −210 mV to −360 mV (**Figure 11C**) and from −410 mV to −610 mV (**Figure 11C**), we could change the relative contributions of heme αII and heme γII to the FTIR difference spectrum and observed significant differences in the spectra of Figure 11C and 11D, notably in the 1700-1550 cm^-1^ region. Although we probably cannot completely separate the contributions from the two low potential hemes, the FTIR spectra were consistent with the dominant contribution of distinct heme species in the two different low-potential windows.

**Figure 11:**
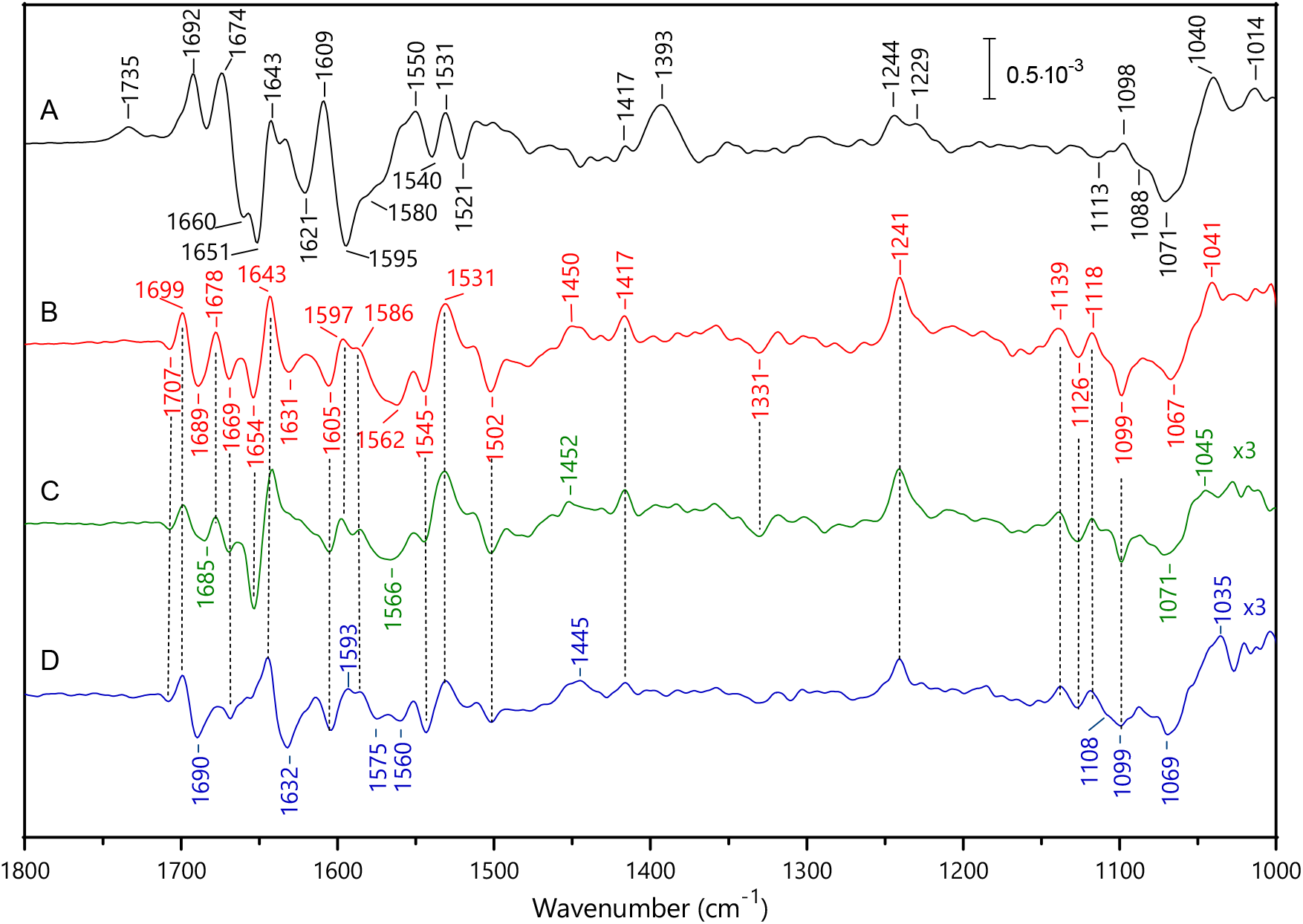
Electrochemistry-induced FTIR difference spectra of ‘Candidatus Kuenenia stuttgartiensis’ hydrazine synthase at pH 8, −210 mV minus +390 mV (**A**), −210 mV minus −560 mV (**B**), −210 minus 360 mV (**C**), and −410 mV minus −610 mV (**D**). Spectra of C and D are enlarged 3 times compared to spectrum B.

Previous thorough investigations of FTIR difference spectra associated with the reduction of protoporphyrin-(methyl)imidazole complexes, 5 coordinated high spin (5c-HS) and 6 coordinated low spin (6c-LS) microperoxidase and *c*-type cytochromes identified redox- and spin-state sensitive IR modes of the heme and axial ligands (21-23). By comparison with these data, positive bands at 1531 cm^-1^, 1417cm^-1^, and 1244-1241 cm^-1^ in the spectra of Figure 11 were assigned to the ν_38_ (CbCb), ν_41_ (CaN), ν_42_ δ(CmH) skeletal heme modes of the reduced heme species (21, 22). Reduced 6c-LS hemes gave rise to an intense ν_42_ δ(CmH) heme mode at 1240-1244 cm^-1^, while this mode was of much lower intensity and broader for a reduced 5c-HS heme (22, 23). The spectrum in Figure 11A significantly differed from the other spectra of Figure 11 by the intensity of the band detected at ≈1242 cm^-1^. This was in line with a heme that is mainly in the 5c-HS form in the reduced state, while the two other hemes had a clear 6c-LS character in the reduced state. Spectral differences were also observed between redox transitions from 6c-LS Imidazole-microperoxidase complexes and 5c-HS microperoxidase in the 1146-1115 cm^-1^ range, where contributions from the ν_44_ (Pyr half ring) heme modes were expected to contribute (22). The spectrum of Figure 11A also clearly differed from the others in this spectral range, in support of its assignment to the HS 5c population of heme γI.

The spectrum recorded for heme γI (**Figure 11A**) showed a positive band at 1734 cm^-1^, in a spectral range where only protonated carboxylic (COOH) groups are expected to contribute. This positive band (i.e. without negative counterpart above 1690 cm^-1^) indicated that heme γI reduction was accompanied by the protonation of a carboxylate group. A negative band at 1580 cm^-1^ is in line with the contribution from a corresponding deprotonated carboxylate group (COO^-^) in the oxidized state (24). In addition, IR modes at 1072/1041 cm^-1^ were assigned to a change in the vibrational contribution of the Tris buffer that corresponded to a deprotonation from the Tris upon heme reduction (23). This suggested that reduction of heme γI was accompanied by a proton uptake from the buffer to a carboxylate group.

The ν(C=O) mode of protonated propionic groups is expected to contribute at 1700–1665 cm^−1^, i.e. at lower frequencies than the corresponding mode of carboxylic groups of Asp or Glu, because of the influence of the heme macrocycle (25, 26). A frequency above 1728 cm^-1^ was proposed, however, for a protonated propionic group of heme b_D_ in dihemic quinol:fumarate reductase (27), but this high ν(C=O) mode frequency was rationalized by a hydrophobic environment of the propionic acid. According to these literature data, the interactions formed by the propionate groups of heme γI in hydrazine synthase did not indicate a possible contribution at 1734 cm^-1^ in Figure 11A. They are both involved in hydrogen bonding interactions with water and/or amino acid side-chains (**Figure 2A**). Therefore, we tentatively assigned the positive band at 1734 cm^-1^ to the redox-coupled protonation of an amino acid carboxylic group. According to the structural environment of heme γI, this group could be Asp168, located at 4.08 Å of the heme iron and stabilizing a water molecule axial ligand of the heme iron (**Figure 2A**). We cannot exclude however, a possible contribution of Glu122, in hydrogen bonding interaction with one of the propionates of heme γI.

Spectrum in Figure 11A showed an intense difference band at 1608/1595 cm^-1^ and a positive band at 1393 cm^-1^ that could correspond to the ν_as_(COO^-^) and ν_s_(COO^-^) modes of a heme propionate group. Indeed, the propionate ν_as_(COO^−^) and ν_s_(COO^-^) modes have been reported at 1620–1540 cm^− 1^, and 1420–1360 cm^− 1^ (26). The high intensity of the band at 1393 cm^-1^ probably points to a specific ionic interaction. For heme γI, one of the propionates forms an interaction with the side chain of Arg 143 (**Figure 2A**) that may be altered upon the heme reduction. Such a change in interaction could also result in changes in the IR mode of the arginine side-chain. The ν_as_(guanidium) vibrational mode was reported at ≈1673 cm^-1^ in solution (24) and at higher frequencies (1680-1695 cm^-1^), upon interaction of the positively charged guanidium group with oxyanions (28). Further investigation, notably using samples in D_2_O, will allow testing this hypothesis and possible contribution of Arg at 1692 or 1674 cm^-1^ in the electrochemistry-induced difference spectrum of heme yI.

As previously mentioned, the FTIR difference spectra shown in Figure 11C and D were typical of those expected for 6c-LS hemes, and presented similarities, concerning the frequencies of the heme skeletal modes. A negative band was observed at 1126 cm^-1^ in these spectra, which corresponded to that previously assigned to the ν(Pyr Half ring) mode in oxidized LS 6c Imidazole-Microperoxidase complex (22). A first difference between these two spectra concerned the IR signature of the axial histidine ligands in the oxidized state. The ν(C_5_-N_1_) ring mode of the histidine axial ligands was identified at 1118 cm^-1^ for the reduced state and as a strong band at 1099 cm^-1^ in Figure 11C for oxidized heme γII, while the corresponding band was more complex with a shoulder at 1108 cm^-1^ in spectrum 11D, containing dominating contributions from heme αII. The frequency of this band depends on the strength of the Fe-Histidine interaction, on the type of coordination to the iron, Nσ or Nπ, and on the protonation state or electronegativity of the histidine imidazole ring (21, 22, 29, 30). The band at 1104 cm^-1^ is typical for a Nσ coordinated neutral histidine side chain, while the band at 1099 cm^-1^ could correspond either to (Nσ coordinated) deprotonated –or highly electronegative-ligand of the heme iron or possibly to a Nπ coordinated neutral side chain (29, 30). Given the lack of hydrogen bonding interactions that could stabilize an imidazolate as seen in the crystal structure, we propose that at least one of the Histidine ligands of heme γII is Nπ coordinated to the heme iron.

Finally, large differences between the FTIR difference spectra recorded upon reduction of heme γII (**Figure 11C**) and heme αII (**Figure 11D**) were observed in the 1700-1600 cm^-1^-region, with strong negative bands at 1690 and 1632 cm^-1^ present only in spectrum of heme αII shown in Figure 11D. This region is complex as peptide groups as well as side chain modes from Arg, Asn or Gln are expected to contribute. Since Heme αII has both propionates involved in ionic interaction with arginine side-chains (**Figure 2D**), it will be interesting to further analyse this region to investigate possible contribution from these interactions.

## Discussion

In this study we combined optical, EPR and FTIR spectroscopy with structural data to attribute redox midpoint potentials and spectral characteristics to the individual hydrazine synthase hemes and gave information about their protein and ligand environment (**Table 1**).

**Table 1:** Spectral characteristics, redox midpoint potential and amino acid residues affected by redox transitions of the different hemes from hydrazine synthase.

| | Heme γII | Heme γI | Heme αII | Heme αI |
| --- | --- | --- | --- | --- |
| VIS | Ox:<br>403 nm<br>Red:<br>420, 553 | Ox:<br>403 nm<br>CT: 620 nm<br>Red:<br>420, 430, 554 nm<br>Mixed HS/LS | Ox:<br>403.5 nm<br>Red:<br>420, 548, 554.5 nm<br>Split alpha band | Ox:<br>398.5 nm<br>CT: 615 nm |
| EPR | $g = 3.46$ | $g = 6$;<br>$g = 2.57/2.27/1.80$<br>Fe-NO as 5<sup>th</sup> ligand<br>$g \approx 2$ ($\Delta G = 16$) | $g = 3.08$ | $g = 6.29/5.65$ |
| $E_m$ (vs SHE) | $\sim -310$ mV | $\sim -30$ mV | $\sim -360$ mV | Irreducible |
| FTIR | COOH, Fe-His | Arg, COO<sup>-</sup>, Fe-His, Asp/Glu, | Arg, COOH, Fe-His | - |

Low spin heme αII had the lowest redox midpoint potential of all hemes in the enzyme and exhibited a spilt alpha band. High spin heme αI could not be reduced and exhibited a rhombic EPR signal conferred by its tyrosine ligation. This heme is proposed to catalyze the comproportionation reaction of hydroxylamine (NH_2_OH) with ammonium. No net electron exchange is involved in this reaction. Su and Chen (31), however, proposed from DFT calculations that the reduced state of the heme would be the lowest energy state in an NH_2_OH-bound intermediate state and should therefore be the active redox state. Our data did not confirm this hypothesis since heme αI remained oxidized in all tested conditions (addition of substrate to the sample (data not shown) did not changed this). We are also confident that the observed signal was not due to an artefact of our purification procedure since it was already present in the crude cell extract and there, too, did not react with reductants. A choice of atoms included in the calculations of Su and Chen (31), probably too restrictive, and an omission to calculate the energies for states M2 and M3 for the oxidized heme may be part of an explanation for these discrepancies. The here presented data may encourage a new round of calculations, with an oxidized heme αI as starting point.

Low spin heme γII showed a redox midpoint potential about 50 mV higher than heme αII, whereas high spin heme γI adopted multiple spin and redox states. The observed redox transitions of γI were higher than the ones for the low spin hemes and occurred around 0 mV. In its reduced state part of the population bound NO.

The γ-subunit is homologous to bacterial di-heme cytochrome *c* peroxidases (bCcP) and MauG proteins. It indeed showed some spectral characteristics reminiscent of these proteins, but also some differences. In bCcP one of the hemes is low spin with a highly anisotropic (HALS) EPR signal, the other one high spin with an axial g=6 signal. In MauG both hemes are low spin with exception of a very small high spin population. In HZS heme γII had a HALS signal, as was the case in bCcP. Heme γI, however, could adopt different spin states as reflected by the EPR and VIS signals. Part of the population exhibited an axial EPR signal at g = 6, like bCcP (32, 33) and the minor high spin population in MauG (34). An additional part of the population of heme γI exhibited low spin characteristics, observed as a main contribution in MauG, giving rise to the g = 2.54/2.27/1.88 signals. We do not yet have proof for the sixth ligand that could confer these spectral characteristics to the heme, but a hydroxyl ion as proposed for bCcP (33) would be a plausible hypothesis from the EPR signature (35). This was supported by the hydrazine synthase structure, where a water was detected in hydrogen bonding distance to the heme iron. The fact that we detected spectral characteristics for a six coordination of heme γI in EPR spectra and in optical spectra precluded that they are due to freezing artefacts as suspected for bCcP.

MauG and HZS both are supposed to catalyse redox reactions involving several electrons at their active site. Whereas for MauG strong redox cooperativity was reported for the two hemes, making them behave as a redox entity, no such observation could be made for HZS. Indeed, for MauG the redox cooperativity translated during a redox titration to a two-step reduction with identical spectral characteristics for both steps (36, 37). This was clearly not the case for HZS where not a two but a three electron transition was proposed to occur at the active site during the NO to NH_2_OH reduction. The here determined redox midpoint potentials of the hemes in the y subunit would allow for reduction of NO to hydroxylamine, which occurs at an overall redox potential of −30 mV. We do not know, however, whether catalysis involves several steps and where the three electrons necessary to complete the reaction stem from. Oxidation of an aromatic amino acid side chain, as observed for KatG (38), a member of the MauG family, might be an option that will be investigated in the future. Irreversible binding of NO to a reduced heme also has to be avoided. Further experiments are necessary to investigate whether NO bound as a fifth ligand to a heme, as observed in this work, is an on-pathway intermediate state, as reported for soluble guanylate cyclase (39) or constitutes a stable off-pathway species, as seen in cytochrome P460 (40).

A further interesting observation concerned the kinetics of heme reduction: heme αII was rapidly reduced by dithionite as expected for a surface exposed heme (**Figure 3, red line**). Heme γII which is exposed to the surface of the protein as well, however, needed minutes to get reduced by dithionite as did heme γI. The sluggish reduction behaviour of the hemes in the γ-subunit, despite their more positive equilibrium redox midpoint potentials, when compared to heme αII, hints to structural and/or ligand environment changes linked to the redox transition. This was strengthened by the variability in spectroscopic signatures and redox midpoint potential observed for heme γI over different redox cycles and between different experiments. Further FTIR experiments in the presence of substrates and in D_2_O buffers might help understanding the protein dynamics surrounding heme γI.

In conclusion, we could identify and attribute distinct spectral properties to the four different hydrazine synthase hemes and determine their redox properties. This data provides the basis to further unravel the reaction mechanism of this unique enzyme.

## Acknowledgements

We would like to thank Guylaine Nuijten and Femke Vermeir for maintenance of the ‘*Candidatus* Kuenenia stuttgartiensis’ enrichment cultures. We would like to thank Frédéric Biaso for helpful discussions about the DFT calculations. Furthermore, we would like to thank Marty Herregraven, Thijs Jacobs and Mark van de Hei from the Technical centre at Radboud University for their technical assistance. We thank the MOSBRI TNA (MOSBRI-2021-33) for providing access to the Aix-Marseille EPR center and FEMS for providing support for FTIR spectroscopy through a research and training grant to WV (FEMS-GO-2021-065). WV and LvN are supported by a NWO Vidi (Vi.Vidi.192.001) to LvN.

